# 5-HT_2C_ receptor perturbation has bidirectional influence over instrumental vigour and restraint

**DOI:** 10.1101/2020.12.13.422360

**Authors:** Oliver Härmson, Laura L. Grima, Marios C. Panayi, Masud Husain, Mark E. Walton

## Abstract

The serotonin (5-HT) system, particularly the 5-HT_2C_ receptor, has consistently been implicated in behavioural control. However, while some studies have focused on the role 5-HT_2C_ receptors plays in regulating motivation to work for reward, others have highlighted its importance in response restraint. To date, it is unclear how 5-HT transmission at this receptor regulates the balance of response invigoration and restraint in anticipation of future reward. In addition, it remains to be established how 5-HT_2C_ receptors gate the influence of internal versus cue-driven processes over reward-guided actions. To elucidate these issues, we investigated the effects of administering the 5-HT_2C_ receptor antagonist SB242084, both systemically and directly into the nucleus accumbens core (NAcC), in rats performing a Go/No-Go task for small or large rewards. The results were compared to administration of *d*-amphetamine into the NAcC, which has previously been shown to promote behavioural activation. Systemic perturbation of 5-HT_2C_ receptors – but crucially not intra-NAcC infusions – consistently boosted rats’ performance and instrumental vigour on Go trials when they were required to act. Concomitantly, systemic administration also reduced their ability to withhold responding for rewards on No-Go trials, particularly late in the holding period. Notably, these effects were often apparent only when the reward on offer was small. By contrast, inducing a hyperdopaminergic state in the NAcC with *d*-amphetamine strongly impaired response restraint on No-Go trials both early and late in the holding period, as well as speeding action initiation. Together, these findings suggest that 5-HT_2C_ receptor transmission, outside the NAcC, shapes the vigour of ongoing goal-directed action as well as the likelihood of responding as a function of expected reward.

## Introduction

The central neurotransmitter serotonin (5-HT) has been implicated in the motivational control of behaviour (Mcelroy et al. 1982; Soubrié 1986; Kulichenko and Pavlenko 2004; Cools et al. 2011). Transmission at the 5-HT_2C_ receptor in particular may play a crucial role in this through the regulation of instrumental vigour – i.e. the energisation of a physical goal-directed response, evident as a change in the speed, amplitude or frequency of movements (Salamone et al. 2007; Dudman and Krakauer 2016). For instance, depletion of central 5-HT or antagonism of the 5-HT_2C_ receptor both speed responding and increase willingness to work for reward (Higgins and Fletcher 2003; Fletcher et al. 2007; Simpson et al. 2011; Bailey et al. 2016, 2018; Browne et al. 2017; Silveira et al. 2020). This has resulted in 5-HT_2C_ receptors being considered a possible target for the treatment of disorders of motivation such as apathy (Higgins et al. 2003; Simpson et al. 2011; Bailey et al. 2016, 2018; Browne and Fletcher 2016; Browne et al. 2017).

However, a largely separate literature has highlighted a key function for intact 5-HT signalling – and again, the 5-HT_2C_ receptor in particular – in enabling appropriate response restraint. Tonic firing of 5-HT neurons increases whilst waiting for reward and decays in the period before an animal ceases to wait (Miyazaki et al. 2011), and depletion of central 5-HT or administration of a 5-HT_2C_ antagonist can also increase inappropriate motor responses in rodents, particularly in anticipation of reward (Winstanley et al. 2004b; Fletcher et al. 2007; Robinson et al. 2008; Agnoli and Carli 2012; Quarta et al. 2012; Adams et al. 2017; Higgins et al. 2020; Silveira et al. 2020). Therefore, a fundamental yet unaddressed question is under what circumstances transmission at 5-HT_2C_ receptors promotes action over inaction and how this might interact with potential future reward.

One possible reason for this lack of clarity is that most studies to date have required animals to work for a constant reward, yet the activity of 5-HT neurons is modulated both by reward magnitude and reward context (Cohen et al. 2015; Li et al. 2016; Matias et al. 2017). A second issue is that the study of action vigour often uses internally guided instrumental paradigms, whilst those investigating the role of serotonin in action restraint have predominantly used stimulus-driven tasks. Such differences are likely to be important as 5-HT has been implicated in gating sensory processing (Petzold et al. 2009), and it has recently been shown that administration of a 5-HT_2C_ receptor antagonist can specifically reduce the influence of cues over decision-making policies (Adams et al. 2017). Therefore, an important open question concerns whether the influence of 5-HT_2C_ in behavioural control depends on the level of reward expectation and if this varies as a function of temporal proximity to reward-predicting cues.

Therefore, to better understand the role of 5-HT_2C_ receptors in shaping the influence of environmental stimuli and reward expectations on action initiation and restraint, we investigated the effect of SB242084 - a functionally selective ligand that is considered to be an antagonist at 5-HT_2C_ receptors (Kennett et al. 1997; Di Matteo et al. 2000; De Deurwaerdere et al. 2004) - on rats’ performance of a Go/No-Go task designed to separate action requirements from the value of acting appropriately (fig 1) (Syed et al. 2016). We predicted that while disruption of 5-HT_2C_ receptors would invigorate instrumental responding on Go trials, it might in tandem impair action restraint for reward on No-Go trials.

**Fig. 1.**
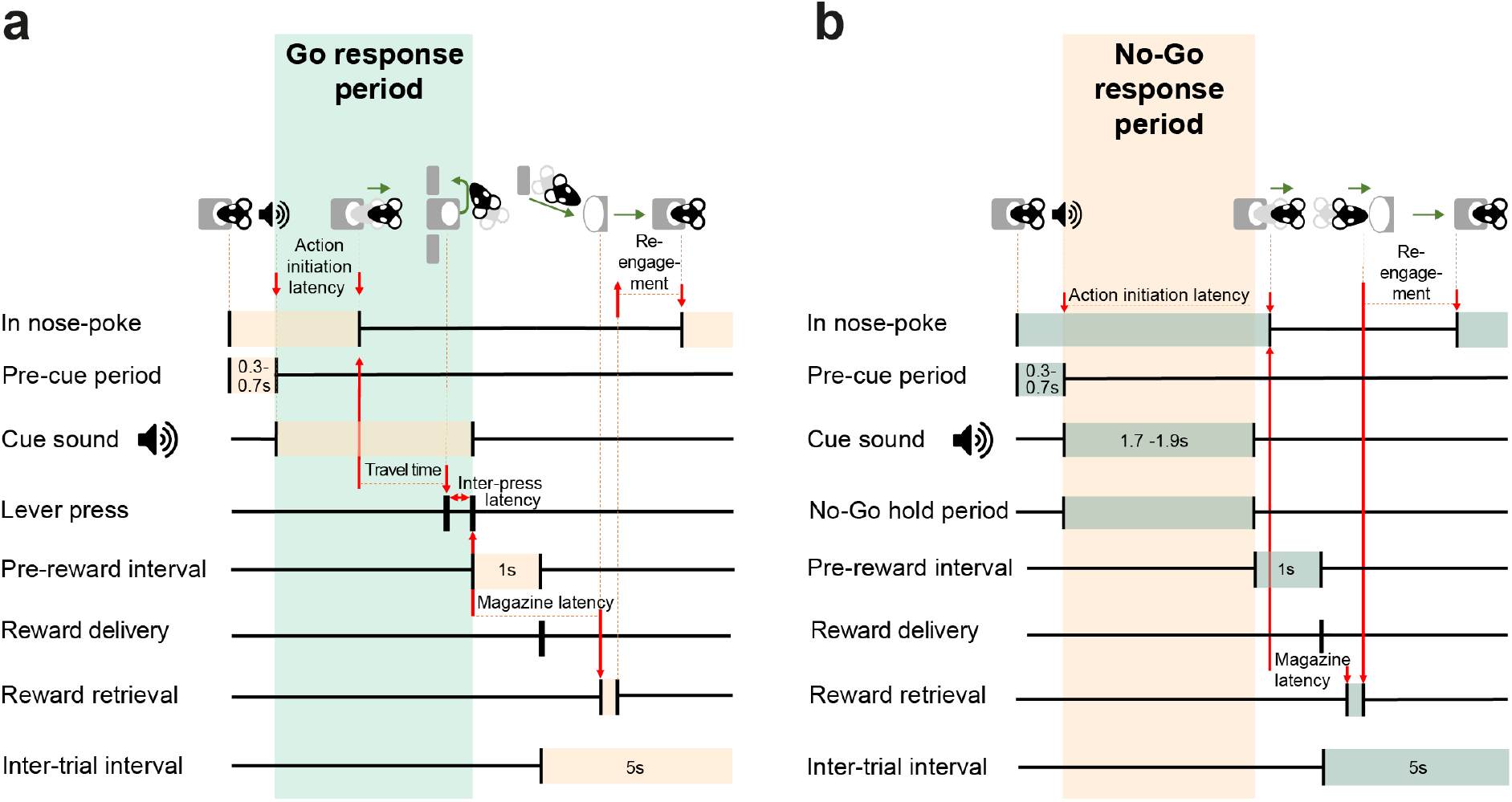
Task epochs in a correctly performed Go (A) and No-Go (B) trial.

After confirming an effect of systemic administration, we further examined whether such effects might be mediated specifically via 5-HT_2C_ transmission in the NAcC. The NAcC is a key site for regulating how motivation translates into action (Robbins and Everitt 1992; Pothuizen et al. 2005; Pattij et al. 2007; du Hoffmann and Nicola 2014; Floresco 2015; Ko and Wanat 2016; Syed et al. 2016). While the majority of studies have focused on how dopamine regulates these processes, the NAcC also receives notable serotonin input (Azmitia and Segal 1978; Vertes 1991) and expresses 5-HT_2C_Rs (Pazos et al. 1985; Eberle-Wang et al. 1997; De Deurwaerdère et al. 2013). Further, disruption of 5-HT_2C_ transmission in the NAcC, but not in medial prefrontal regions, has been shown to increase premature responding on the 5-choice serial reaction time task (Robinson et al. 2008). Therefore, we compared the effects of SB242084 into the NAcC with administration of d-amphetamine, under the hypothesis that both drugs might weaken action restraint for reward.

## Methods

### Subjects

All procedures were carried out in accordance with the UK Animals (Scientific Procedures) Act (1986) and its associated guidelines. 26 group-housed male Sprague-Dawley rats (Envigo, U.K.), aged ~2 months at the beginning of training, were used. Animals were maintained on a twelve-hour light/dark cycle (lights on 07.00). During testing, rats were food restricted to maintain ~85-90% of their free-feeding weight. Water was available *ad libitum* in their home cages. For the NAcC infusion studies, animals were implanted with bilateral guide cannulae (Plastics One) 1.5mm above the target site of the NAcC (AP:+1.4mm, ML:±1.7mm, DV:−6.0mm from skull surface) under isoflurane anaesthesia and secured with dental acrylic.

### Behavioural task

#### Apparatus

Testing was carried out in operant chambers (30.5 x 24.1 x 29.2 cm; Med Associates) (fig. 2a). Each chamber was housed within a sound-attenuating cabinet ventilated with a fan, which provided a constant background noise of ~59 dB. Each chamber contained two retractable levers 9.5 cm of either side of a central nose-poke, which was fitted with an infrared beam signalling when animals entered the receptacle. The wall opposite contained a food magazine into which 45 mg sucrose pellets (Sandown Scientific, UK) could be dispensed. Each chamber was also fitted with a house-light, and a speaker for delivering auditory stimuli.

**Fig. 2.**
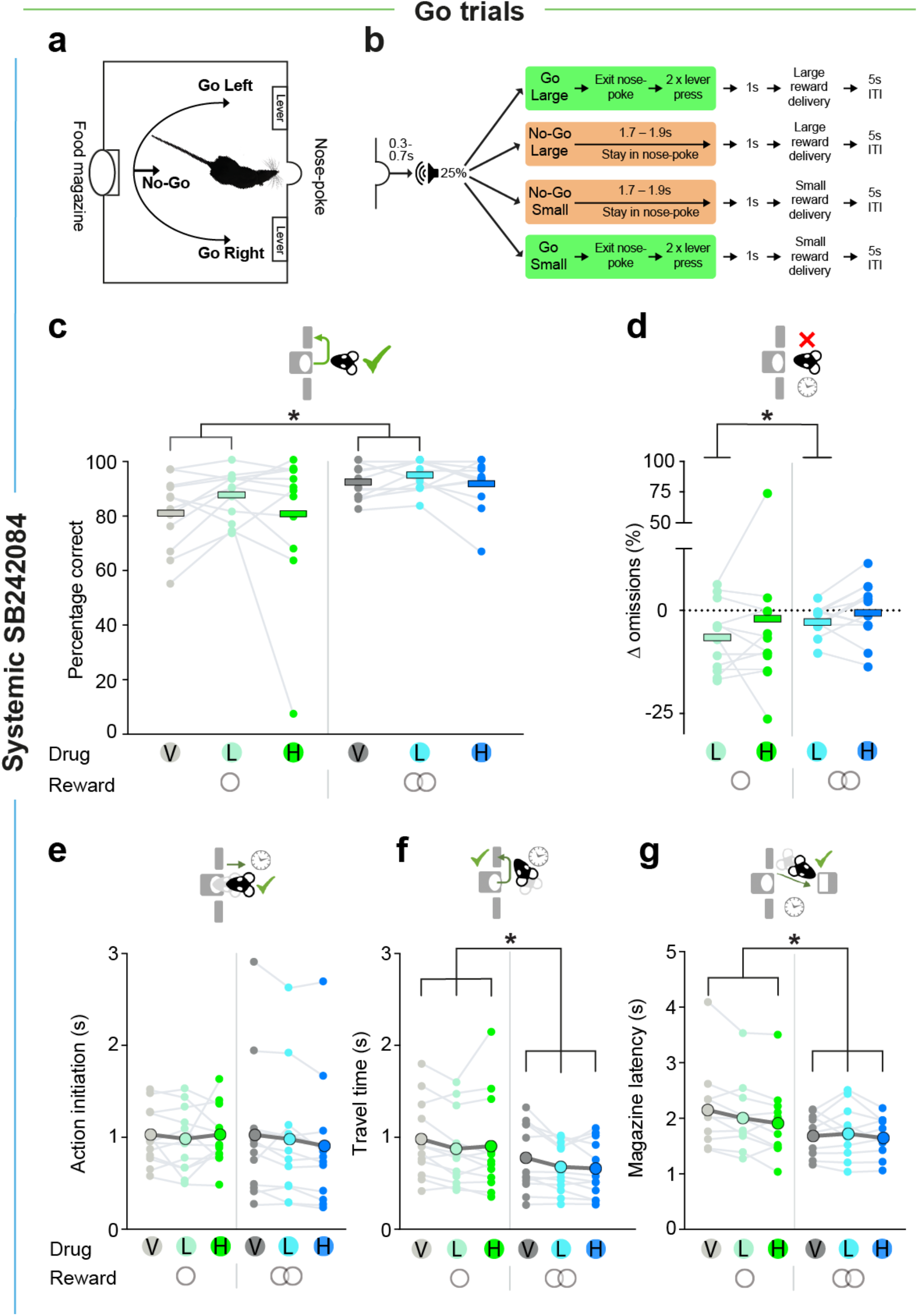
Behavioural setup and the effects of systemic SB242084 on Go trial performance and latencies (n=12) **A, B.** Schematic of the operant box (A) and the trial types (B). **C.** Percentage correct responses on small and large reward Go trials. Pairwise comparisons: vehicle v low dose, p=.004; all other comparisons, p > .166. **D.** Change in percentage of Go trial omissions on small and large reward trials, calculated as the difference between drug and vehicle treatments. Pairwise comparisons: vehicle v low dose, p=.002; all other comparisons, p > .306. Data in c-d are depicted as treatment means (thick bars), superimposed by individual subject data (dots and grey lines). **E.** Action initiation latency on successful small and large reward Go trials. **F.** Travel time on successful small and large reward Go trials. Pairwise comparisons: low dose v vehicle, p=.001; high dose v vehicle, p=.017. **G.** Magazine latency on successful small and large reward Go trials. Pairwise comparisons: vehicle v low dose: p=.187; vehicle v high dose, p=.008. Data in **E-G** are shown as means (large coloured circles), superimposed by individual rat data (lines). *p < .05

#### Paradigm

The task design is shown in figure 2a-b and figure 1. The task required animals to use an auditory cue to guide whether to initiate a response and press either the left or right lever (Go Left/Right) or to withhold responding (No-Go) to gain either a small or large reward. A trial was initiated when the rat voluntarily entered and stayed in a central nose-poke for 0.3-0.7s s. This triggered the presentation of one of four auditory cues, which signalled (a) the action requirement (Go Left/Right or No-Go) and (b) the reward for a correctly performed trial (Small or Large, respectively 1 or 2 sucrose pellets) (fig.2b). Go trials required animals to make two presses on the correct lever within 5 s of cue onset (fig.2b, fig.1a). On No-Go trials, animals were required to remain in the nose-poke for the No-Go hold period (fig. 1b). While No-Go trials always posed the same requirement regardless of the promised reward, the left/right mapping on Go trials was consistently associated with a specific reward size (small/large), with the side and reward associations counterbalanced across the cohort. This allowed us to independently manipulate action requirement and reward expectation, and additionally to assess how this influenced the direction of motor responses on Go trials. Successful trials caused reward to be delivered to a food magazine on the opposite wall of the chamber.

#### Training

To teach rats to respond or withhold actions to go and no-go cues, animals were trained as described by (Syed et al. 2016). Briefly, animals were first habituated to the operant chamber and learned to retrieve pellet rewards from the food magazine tray. Rats then commenced training initially exclusively with the No-Go trial type. Over several sessions, they were gradually trained to make and hold a response in the central nose poke, such that on No-Go trials they were eventually able to withhold responding during a 0.3-0.7 s pre-cue nose-poke hold period and a subsequent No-Go hold period (Cohort 1, systemic SB242084: 1.7-1.9s; Cohort 2, local SB242084, *d*-amphetamine and systemic SB242084 replication: 1.5-1.7s) to gain reward (1 or 2 sucrose pellets respectively for ‘No-Go Small’ or ‘No-Go Large’ trials). The cue was either a tone, buzz, white noise, or clicker, counterbalanced across animals (each ~74 dB). A premature head exit caused the house light to be illuminated as the animal exited the nose poke for the duration of a 5s time-out, after which the house light turned off and a standard 5s inter-trial interval (ITI) commenced.

Once 60% of No-Go Small and Large trials were performed correctly, rats were next trained exclusively on the Go trials. Here, after a 0.3-0.7 s central nose-poke, one of the two remaining auditory cues would sound, one requiring a left lever press and the other a right lever press (side counterbalanced across animals). Correct responding on a particular lever was associated throughout testing with either a single sucrose pellet (‘Go Small’) or two sucrose pellets (‘Go Large’). An incorrect lever press would result in the house light switching on for a 5s time-out period, followed by a 5s ITI. During training, an error-correction procedure was used so that the next trial after an error would always be of the same cue/trial-type with the wrong lever withdrawn. Once a criterion of 60% successful Go responses was reached, interleaved No-Go and Go trials were introduced, each with a 25% probability (other than correction trials).

Once an average 60% success rate on all four trial types in a session was achieved, the rats moved onto the full version of the task. Here, error correction trials were removed. Further, the number of necessary lever presses on Go trials was increased to two. This ensured that the interval between cue onset to reward delivery was similar between Go and No-Go trials. Reward delivery was delayed for 1 s after the successful completion of a trial. Similarly, the error signal (the house-light being illuminated) was also delayed for 1 s following an erroneous response. Throughout training and testing, a session ended after the animals had either earned 100 rewards or had spent 60 min in a session, whichever came first (although rats always met the former criterion during test sessions in the current study). Typically, rats took 1-1.5 months to train to reach stable baseline performance and each rat in the current study had performed the full task at least 10 times before undergoing drug testing sessions. In between every drug testing session, rats underwent at least one training session without drug manipulations to reestablish baseline performance, where error correction trials were reintroduced, but all other parameters were kept constant.

### Pharmacological challenges

Drugs were administered in a within-subjects regular Latin square design. The local infusion sessions were performed following the full completion of an experiment to assess the effect of intra-NAc infusions of D1 agonist or antagonist drugs on task performance (data to be reported separately) and after returning to stable baseline levels of performance. This meant that, prior to the start of local infusion experiments reported here, rats had received 6-7 NAcC infusion (plus one mock infusion session where no substance was injected). Animals always performed at least one behavioural session without injection or infusion in between each drug manipulation to reestablish baseline performance and rule out lasting effects of drugs.

#### Systemic pharmacology

SB242084 (6-Chloro-2,3-dihydro-5-methyl-*N*-[6-[(2-methyl-3-pyridinyl)oxy]-3-pyridinyl]-1*H*-indole-1-carboxyamide dihydrochloride, Tocris) was dissolved in 25 mM citric acid in 8% w/v cyclodextrine in distilled water, and the pH adjusted to 6-7 using 5 M NaOH. Doses for systemic administration match those used in previous studies to have demonstrated increases in impulsive responding (Winstanley et al. 2004b, a; Fletcher et al. 2007; Silveira et al. 2020) and instrumental vigour (Higgins et al. 2003). Stock solutions of the vehicle (25 mM citric acid, 8% w/v cyclodextrine in distilled water), SB242084 0.1 mg/ml (referred to in the results’ section as “Low” dose) and SB242084 0.5 mg/ml (“High” dose) were prepared and aliquoted before being frozen at −20°C. Concentrations of both drugs were calculated as the salt. On each experimental day, one aliquot of each drug was defrosted. Systemic injections of drug or vehicle, containing 25 mM citric acid and 8% w/v cyclodextrine in distilled water, were given intraperitoneally in a volume of 1 ml/kg. Injections were given 20 minutes prior to testing.

#### Local infusions

SB242084 made up as described above. Doses were chosen based on a prior report showing a consistent and dose-dependent increase in impulsive responding on the 5CSRTT (Robinson et al. 2008). Stock solutions (Vehicle, SB242084 0.2 μg/μl, 1.0 μg/μl) were prepared and aliquoted before being frozen at −20°C. Concentrations of both drugs were calculated as the salt. On each experimental day, one aliquot of each drug was defrosted. D-amphetamine ((+)-α-Methylphenethylamine hemisulfate, Sigma-Aldrich) was dissolved in 0.9% NaCl solution to reach a concentration of 10 μg/μl). This was aliquoted and frozen at −20°C. On each experimental day, one aliquot of the drug was defrosted.

### Surgical procedures

Rats were anaesthetised using inhaled isoflurane (4% in O2 for induction and 1.5% for maintenance) and given buprenorphine (Vetergesic, 0.03 mg/kg) and meloxicam (Metacam, 2 mg/kg) at the start of surgery. Once animals were secured in a stereotaxic frame and their scalp shaved and cleaned with dilute hibiscrub and 70% alcohol, a local anaesthetic (bupivicaine, 2 mg/kg) was administered to the area. The skull was then exposed and craniotomies made for implantation of bilateral guide cannulae (Plastics One, UK), consisting of an 8mm plastic pedestal holding together two 26 gauge metal cannulae with a centre-to-centre distance of 3.4mm and a length of 7.5mm. The cannulae were implanted 1.5mm above the target site of the NAcC, at AP +1.4mm, ML ±1.7mm, DV −6.0mm from the surface of the skull and relative to bregma (Franklin & Paxinos, 2007). 4 anchoring screws (Precision Technology Supplies) were also implanted and dental acrylic applied to secure the cannulae to the skull. At the end of surgery, dummy cannulae of the same length as the guide cannulae were inserted to ensure patency and a dustcap was secured onto the pedestal to secure the dummy. Following surgery, animals were again administered buprenorphine (0.03 mg/kg) and meloxicam (2 mg/kg), and were thereafter given meloxicam for up to 3 days post-surgery.

### Local infusion procedure

Animals were first habituated to manipulation of their implants by being lightly restrained and having the dummy cannulae removed and a fresh set reinserted. Re-training of the animals commenced approximately 2 weeks after surgery. All animals returned back to criterion performance (≥60% success rate on each trial type) before drug infusions commenced. During infusions, the rats were gently restrained whilst the dummy cannulae were removed and the 33 gauge bilateral infusion cannulae, at a length of 9mm, were inserted into the NAcC. 0.5 μl of vehicle or drug solution was injected per hemisphere at a rate of 0.25 μl/min. The infusion cannulae were left in place for 2 minutes after the cessation of the infusion to allow diffusion of solution from the cannulae. Next, the infusion cannulae were removed, the dummy cannulae and dust cap replaced, and the rats were returned to their home cage for 10 minutes to reduce possible effects of injection stress on performance. Testing then commenced.

### Histology

At the end of data collection, animals were deeply anaesthetised with sodium pentobarbitone (200 mg/kg, i.p. injection). They were then transcardially perfused with 0.9% saline followed by a 10% formalin solution (vol/vol). The brains were kept in 10% formalin solution until being sectioned. Brains were sectioned into 60 μm-thick coronal sections using a vibratome (Leica). The sections were then stained with cresyl violet (Sigma Aldrich) before being mounted in DePeX mounting medium onto 1.5% gelatin-coated slides and enclosed with coverslips.

### Data analysis

Latency and performance measures of interest are summarized in table 1.

**Table 1.**
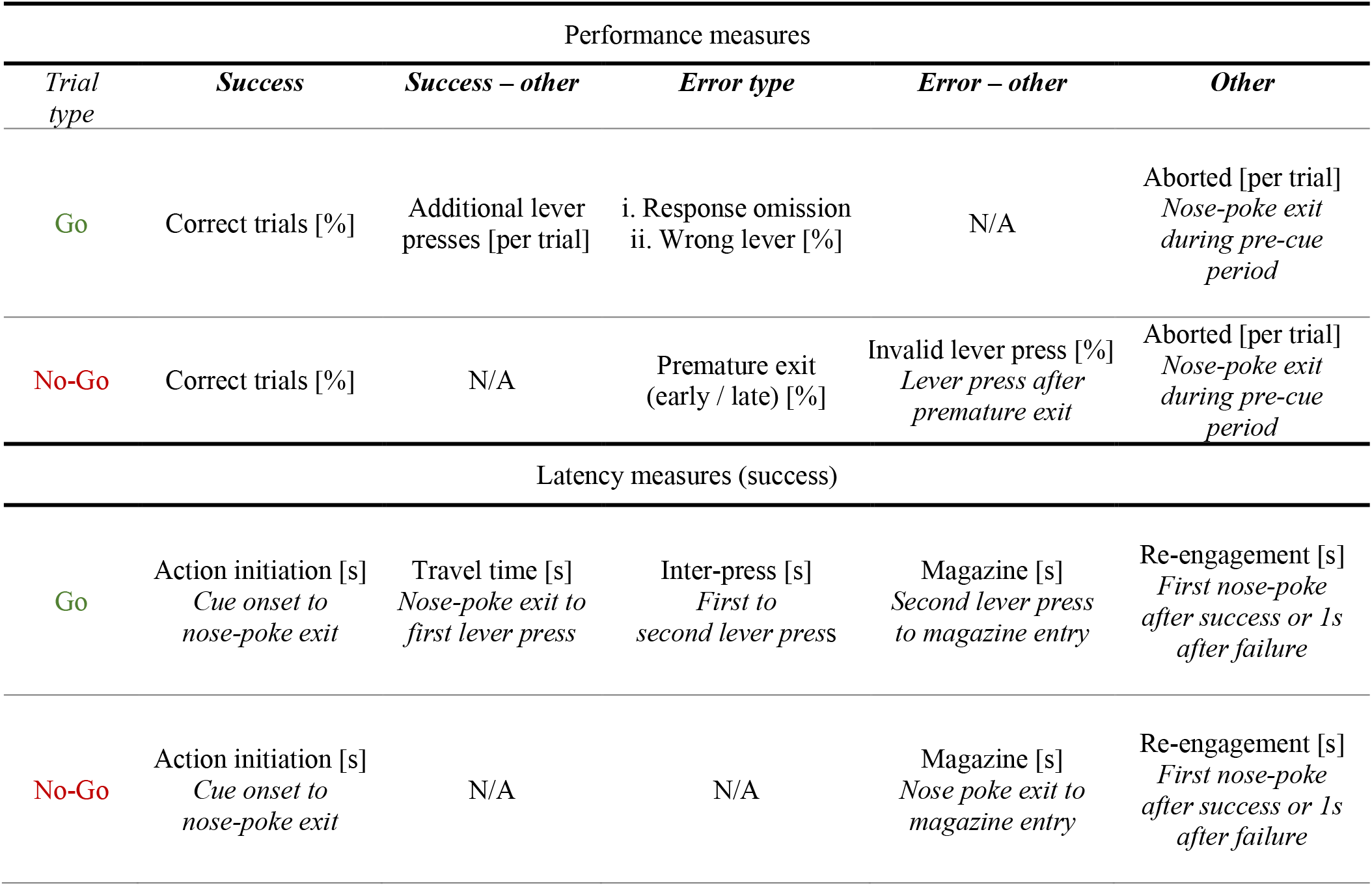
Overview of the behavioural variables of interest within each trial. Further details can be found in Fig. 1. ‘N/A’, not applicable.

#### Measures of performance

The primary measure of performance was **Go** and **No-Go trial success**, measured as percentage of correctly performed small or large reward Go or No-Go trials respectively. On Go trials, we considered two types of errors: (i) no response on either lever within 5 seconds (‘**Response Omission**’ Go) or (ii) incorrect lever response (‘**Wrong Lever**’ Go). No-Go trials were considered to be erroneous if rats left the nose poke prematurely before the end of the holding period while the cue was still playing. We reasoned that premature responses could result from either a failure to inhibit fast cue-driven responses, or from a failure to wait for the appropriate time period before initiating a response. We anticipated that the former process would lead to premature responses clustered near cue presentation, while the latter would manifest in No-Go failures nearer the end of the holding period. In support of this distinction, we had previously observed that stimulation of dopamine receptors in this task selectively increased cue-elicited No-Go errors (Grima et al. 2021). Therefore, instances of premature action initiation were quantified within the first and second halves of the No-Go hold period (‘early’ and ‘late’ epochs, < 0.95 s and ≥ 0.95s or < 0.85s and ≥ 0.85s for the 1st and 2nd cohort, respectively).

Additionally, there were three other behaviours that animals could exhibit that reflected possible changes in performance. First, after exiting the nose poke prematurely on an unsuccessful No-Go trial, animals sometimes pressed a lever. If this occurred during a period equivalent to the minimum No-Go hold interval, these instances were labelled as ‘**invalid lever press**’ trials and quantified. Second, the number of **presses on the correct lever** preceding food magazine entry was quantified in Go trials, as rats sometimes continued to press beyond the required number to gain reward. Third, occasionally rats exited the nose poke prematurely during the pre-cue period and thereby failed to initiate a new trial. These instances were labelled ‘**aborted trials**’.

#### Measures of latency

Trial windows of interest are summarised in Figure 1. **Action initiation latency** was measured as the time from cue onset to exit of the nose poke in both Go and No-Go trials. Following on from the planned analyses of the patterns of No-Go errors, we conducted additional exploratory analyses on successful No-Go trials where the nose poke exits were classified as falling within the pre-reward (between cue offset and reward delivery later) or within the post-reward interval. We analysed observations within the first 0.8s of each of these intervals (instead of the full 1s), as the jitter of the No-Go hold duration sometimes resulted in the remaining 0.2s falling outside of the time interval of interest. Similar to our conceptualisation of the No-Go errors, we reasoned that responses clustered just after No-Go cue offset in the pre-reward period or just after reward delivery in the post-reward interval could be considered as cue-driven responses. **Travel time** in Go trials was measured as the time from nose poke head exit to 1^st^ press on the *appropriate* lever. **Inter-press latency** was measured as the time interval between the 1^st^ and 2^nd^ press on the appropriate lever. **Magazine latency** was defined as the time interval between completion of a successful No-Go response or the Go response and entry to the food magazine. Changes in within-trial response latencies were used as proxies for changes in animals’ instrumental vigour. **Reengagement latency** was defined as the time taken to re-enter the nose-poke either after leaving the food magazine in successful trials or 1s following the registration of an error on unsuccessful trials.

#### Intra-NAcC d-amphetamine

In order to determine whether any of the effects in the local SB242084 study might have been masked by tissue damage around the tips of the cannulae, we also carried out intra-NAcC infusions of d-amphetamine in this cohort, which has previously been shown to increase behavioural activation and premature responding (Cole and Robbins 1987; West et al. 1998). This meant that, as well as analysing the effect of a hyperdopaminergic state in NAcC on task performance and latencies, we could also use two behavioural measures that were predicted to be strongly influenced by this drug in the majority of animals – *No-Go accuracy* and *Go trial action initiation latency* – as a potential marker for the continued viability of the brain tissue around the cannulae. Specifically, if the effect of *d*-amphetamine on these measures within a subject did not exceed the cohort mean *d*-amphetamine-induced change by ≥50% in *at least one* of these behavioural outcomes, that animal was categorized as potentially having functionally significant tissue damage around the cannulae tips.

### Statistical approaches

All data were analysed using Matlab (Mathworks), SPSS (IBM) and R (The R Foundation). Performance and time measures were mainly analysed using repeated-measures ANOVAs with drug dose and reward size as the within-subject factors, unless specified otherwise. As we have described the effects of reward on task performance in a detail in a separate manuscript (Grima et al. 2021), the primary focus here was on effects of and interaction with drug. Any significant main effects of reward are, however, reported in the figure legends. Behavioural measures not reported in the main text are documented in Supplementary Tables 1-3 (1: Systemic SB242084; 2: Intra-NAcC SB242084; 3: Intra-NAcC d-amphetamine). In the d-amphetamine infusion experiment, we used sessions where saline was locally infused into NAcC from a parallel dopamine receptor pharmacology experiment in the same animals as vehicle data for within-subjects analysis. In the systemic SB242084 replication experiment, an additional within-subject factor ‘training experience’ was specified, due to the inclusion of 2 separate replications of systemic SB242084 administration, one prior to cannulation surgery and one after local infusion experiments had been completed. Percentages of nose-poke exits in the early and late parts of the No-Go hold period were compared using the chi-squared test of contingency on the group level, as the low number of such observations resulted in skewed residual distributions in ANOVAs. Whenever there was a significant main effect of drug or an interaction with a task variable of interest, test results were reported in the main text or figure legends and *post hoc* comparisons across levels of that effect were carried out. The influence of outliers on the ANOVA results was minimized by excluding any subject on the basis of absolute standardised residual values bigger than 3 from that analysis. A p-value less than 0.05 was considered significant.

## Results

### Systemic SB242084 increases accuracy and instrumental drive on Go trials

We first examined how SB242084 influenced action selection on Go trials, where rats were required to initiate an action and press the appropriate lever twice for small or large reward. In vehicle sessions, rats’ success rate was on average >80% on both trial types.

A first analysis did not find any significant effects of the drug on Go performance (all F<1.49, p>.248). However, as prior research has reported SB242084 to have non-linear effects on performance with increasing drug doses (Fletcher et al. 2007), we sought to test for non-linearities by running within-subject contrasts. This revealed that the 5-HT_2C_ antagonist caused an overall *improvement* in performance for both low and high reward trials selectively at the low dose, where 11 of the 12 rats showed greater than average success rates compared to vehicle (quadratic effect of drug: F_1,11_=11.26, p=.006; drug X reward interaction: F_2,22_=0.29, p=.789; fig.2c). Further analyses showed the improvement in performance on the low dose was caused by a decrease in lever press omissions (quadratic effect of drug: F_1,11_=7.73, p=.018; fig. 2d), with no corresponding change in the frequency of incorrect lever response trials (all F<1.43, p>.261; table S1).

We next examined whether administration of the drug altered motor responses within Go trials. While there was no reliable change in latency to initiate action, (all F<1.65, p>.216; fig.2e), administration of either dose of the drug reduced the travel time to the correct lever (main effect of drug: F_2,22_=7.15, p=.004; fig.2f) and decreased the inter-press latency (main effect of drug: F_2,20_=4.41, p=.026; pairwise comparisons: high dose v vehicle, p = .004, all other ps>.199; table S1) as well as the magazine latency (main effect of drug: F_2,18_=6.71, p=.007; fig.2g). Despite this invigoration, the ligand did not change the number of lever presses made on the correct lever (all main effects and interactions: Fs<1.91, ps>.176; table S1). These findings suggest that systemic SB242084 promotes engagement with the task and invigorates ongoing motor responses for rewards, and, at a low dose, can do so without compromise to instrumental precision.

### Systemic SB242084 increases impulsive action on No-Go trials

We next considered the effect of SB242084 on rats’ ability to withhold action for reward by comparing the proportion of correct small and large reward No-Go trials in each of the drug administration conditions. On vehicle, animals successfully withheld responding for small and large rewards on average on >84% of No-Go trials. However, in contrast to Go trials, administration of either dose of the drug *impaired* performance when the prospective future reward was small (drug x reward interaction: F_2,22_=5.18, p=.014; fig. 3a). This was not caused by a general inability to withhold action as not only did the ligand have no effect on performance when the large reward was on offer, it also did not change the number of aborted trials, when the rats failed to sustain the precue nose poke required to initiate a new trial (no main effect or interactions with drug: F<0.93, p>.409; table S1). Further, the magnitude of the impairment did not reliably correlate with each animal’s improvement in Go trial success for either dose of the drug (all −0.30 > Spearman r > −0.08, all p > 0.34).

**Fig. 3.**
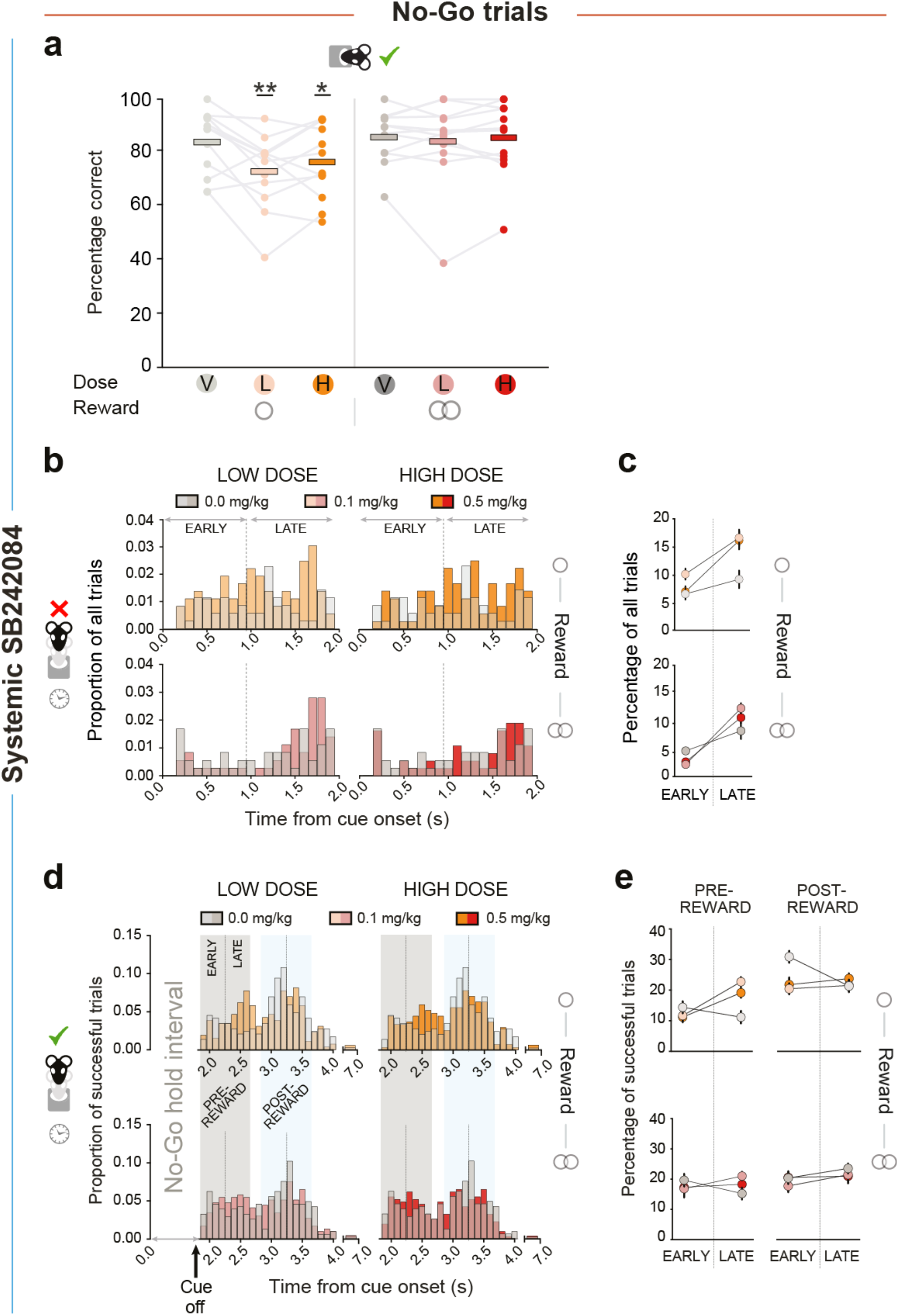
The effects of systemic SB242084 on No-Go trial accuracy and nose poke head exit timing (n=12) **A.** Percentage of correct responses on small and large reward No-Go trials performed per drug session. Data are shown as mean (thick bars), superimposed by individual subject data (dots and grey lines). Pairwise comparisons: low dose and high dose v. vehicle on small reward trials: p=.005 and p=.039, respectively; all other, p > .236. *p < .05, **p < .01 denoted as comparisons against the vehicle group on small reward trials. Main effect of reward: F_1, 11_ = 4.98, p = 0.047. **B,C.** Timing of premature nose poke exits on unsuccessful small (upper panel) and large (lower panel) reward No-Go trials. Distributions are normalised to overall numbers of small or large reward No-Go trials. Data are shown in distributions of 100ms bins (B) or divided into ‘early’ and ‘late’ epochs of the No-Go hold requirement (C). **D,E.** Timing of action initiation on successful small (upper panel) and large (lower panel) reward No-Go trials. Highlighted are the pre- and post-reward intervals (grey and light blue shading, respectively), each of which consist of an early and a late epoch. Data are shown in distributions of 100ms bins (D) or divided into ‘early’ and ‘late’ epochs of either the pre-reward or post-reward period (E). Distributions are normalised to numbers of correct small or large reward No-Go trials. Pairwise comparisons: low dose v. vehicle during the ‘early’ epochs p=.018; low dose v. vehicle during the ‘late’ epochs: p=.001; all other p > .070. Data in C and E are depicted as mean ± SEM, normalized to the within-subject variance across small and large reward trials either just within the No-Go hold period (C) or within the pre-or post-reward interval (E).

We reasoned that successful action restraint during the No-Go hold interval relies, first, on inhibition of reactive responses - triggered by the onset of the cue – followed, second, by appropriate control of anticipatory responses targeted at reward retrieval. To test this more directly, we quantified premature head exits in either the ‘early’ or ‘late’ epoch of the No-Go hold period of error trials (fig.3b). This revealed that both doses of the ligand promoted erroneous ‘late’ over ‘early’ head exits, again only when the rats were anticipating a small reward (small reward: vehicle v low dose: χ^2^(2)=9.16, p=.010; vehicle v high dose: χ^2^(2)=6.86, p=.032; large reward: all χ^2^ <5.26, p>.072; fig.3c).

We also investigated whether SB242084 affected action initiation latencies following the successful completion of the No-Go waiting requirement. Although there were no significant differences in correct No-Go trial hold durations on and off the ligand (all F<0.83, p>.451), based on the patterns of No-Go errors we predicted the ligand might have also altered the balance of fast and slow head exit timings with respect to salient task events (cue offset, signalling the end of the No-Go period, and reward delivery 1s later). We therefore examined the distribution of head exit latencies on correct No-Go trials within the pre-reward interval (i.e., in the 1s between No-Go cue offset and reward delivery) as well an equivalent post-reward interval, both split into ‘early’ and ‘late’ epochs. As can be observed in figure 3d, rats given vehicle injections on average had the highest likelihood of leaving in the early part of each interval. However, this pattern switched such that the rats given the ligand – particularly at the lower dose – became more likely to leave in the late part of each interval (early-late X drug interaction: F_2,22_=5.81, p=.009; fig 3e). Similar to Go trials, once an action had been initiated, the drug dose-dependently reduced magazine latencies (main effect of drug, F_2,18_=7.51, p=.004; vehicle v low dose, p=.006; vehicle v high dose, p=.023; table S1).

In summary, SB242084 selectively impaired animals’ ability to withhold responding when the reward on offer was small. This reflected a significant increase in inappropriately leaving the nose-poke port late in the hold period without a concomitant increase in early cue-driven responses. This shift from early to late responses persisted on successful No-Go trials, both in the pre-reward and post-reward interval.

### Effects of 5-HT_2C_ receptor manipulation on action restraint / instrumental drive not localisable to NAcC

It has previously been suggested that 5-HT_2C_ receptors in the NAcC are important for incentive motivation and inhibitory control (Robinson et al. 2008). Therefore, to determine whether the attenuation of goal-directed inhibition and invigoration of reward-related lever pressing we observed after systemic injections was dependent on 5-HT_2C_ receptors in the NAcC, we tested a second cohort of animals on the Go/No-Go task following local infusion of either vehicle, 0.1μg or 0.5μg per hemisphere of SB242084. Of the 14 implanted animals, one was excluded for having misplaced cannulae and two others did not complete all testing sessions, resulting in an n of 11 rats (fig.4a).

**Fig. 4.**
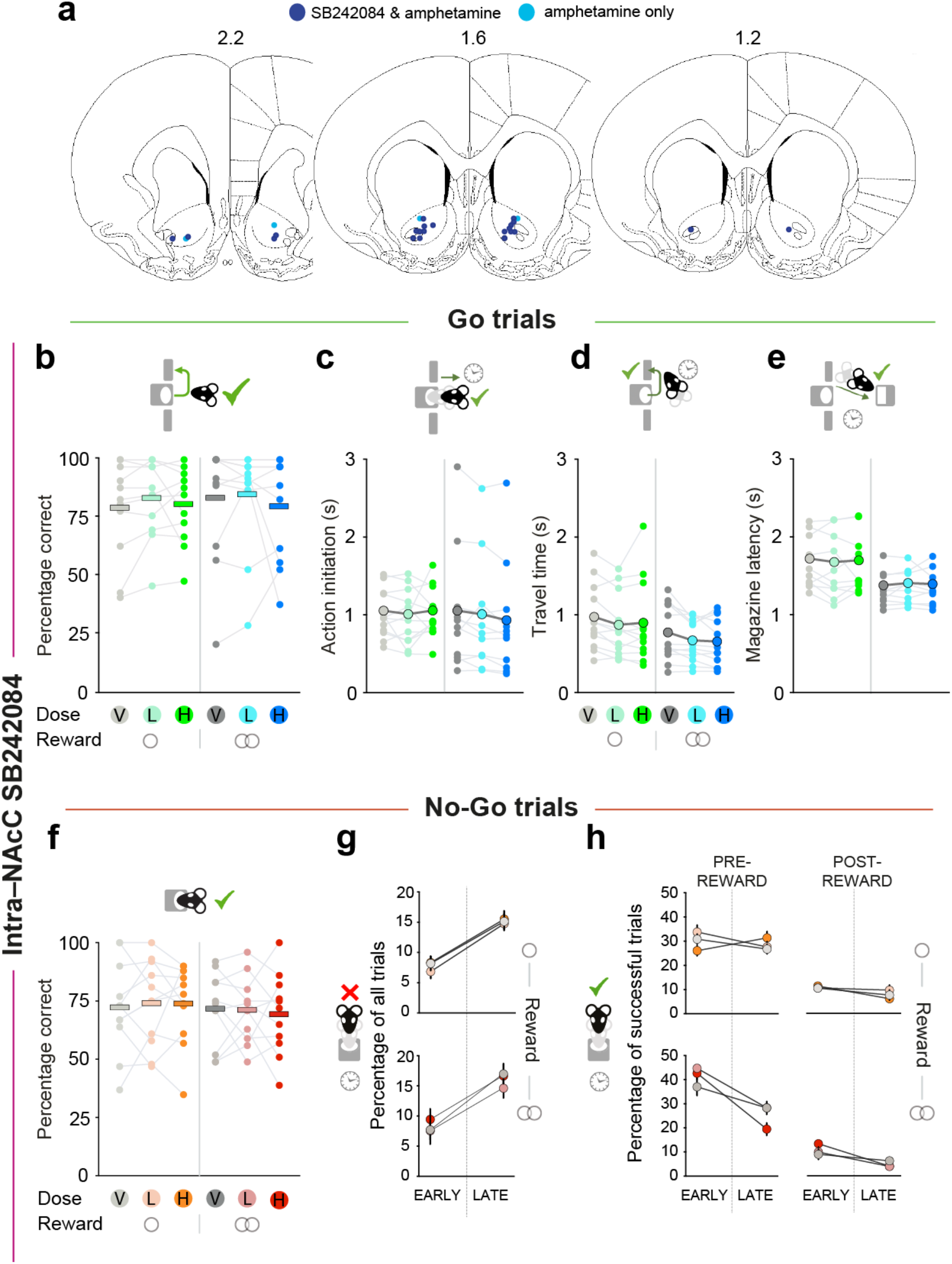
Effects of intra-NAcC infusions of SB242084 on Go/No-Go performance and latencies. **A.** The locations of the tips of the guide cannulae depicted on horizontal section schematics adapted from Paxinos and Watson (2009). **B,F.** Percentage of correct responses for small and large reward trials performed per drug session in Go (B) and No-Go (F) trials. Data are shown as means (horizontal bars), superimposed by individual subject data (dots and light grey lines). **C-E.** Effect of intra-NAcC vehicle or SB242084 on action initiation (C), travel time (D) and magazine latency on Go trials (E). Data are depicted as means (large dots), superimposed by individual subject data (thin grey lines and small dots). Travel time: main effect of reward: F_1, 10_ = 5.95, p = 0.035; magazine latency: main effect of reward: F_1,9_ = 10.10, p = 0.011. **G.** Effect of intra-NAcC vehicle or SB242084 on the timing of premature nose poke exits, taking place in either the early or late epochs of unsuccessful No-Go trials (small or large reward, upper and lower panel, respectively). **H.** Effect of intra-NAcC vehicle or SB242084 on timing of nose-poke head exits in the early or ‘late’ epochs of the pre-reward interval and post-reward interval of successful No-Go trials. Data in G and H are depicted as mean ± SEM, normalized to the within-subject variance across small and large reward trials either just within the No-Go hold period (G) or within the pre-or post-reward interval (H).

In contrast to the effects of systemic administration, and against our expectations, there were no reliable effects of intra-NAcC SB242084 compared to vehicle on either Go or No-Go trial success rate or latencies (all F<0.97, p>.395; fig.4b-h; table S2). This remained the case even when limiting analyses to animals that showed a change following d-amphetamine infusions (see next section).

### Amphetamine in NAcC impairs action restraint but does not improve instrumental performance

The lack of effect of intra-NAcC infusions of the ligand appears to suggest no direct role for 5-HT_2C_ receptors for performance on this task. However, it is also possible that it reflects a failure of the drug to reach viable brain tissue or a difference in the behavioural strategy of this cohort of animals from the group that were used for the systemic manipulation.

To rule out the former explanation, and also to provide a direct comparison with a dopaminergic manipulation, we next examined the effect of intra-NAcC infusions of 5μg per hemisphere of d-amphetamine in the same cohort of animals (n=13). Infusions of d-amphetamine did not influence Go trial success rate (F < 0.24, p > .630) (fig.5a). Nonetheless, the drug substantially and selectively speeded action initiation on successful Go trials (main effect of drug: F_1,12_=12.74 p=.004; fig.5b).

**Fig. 5.**
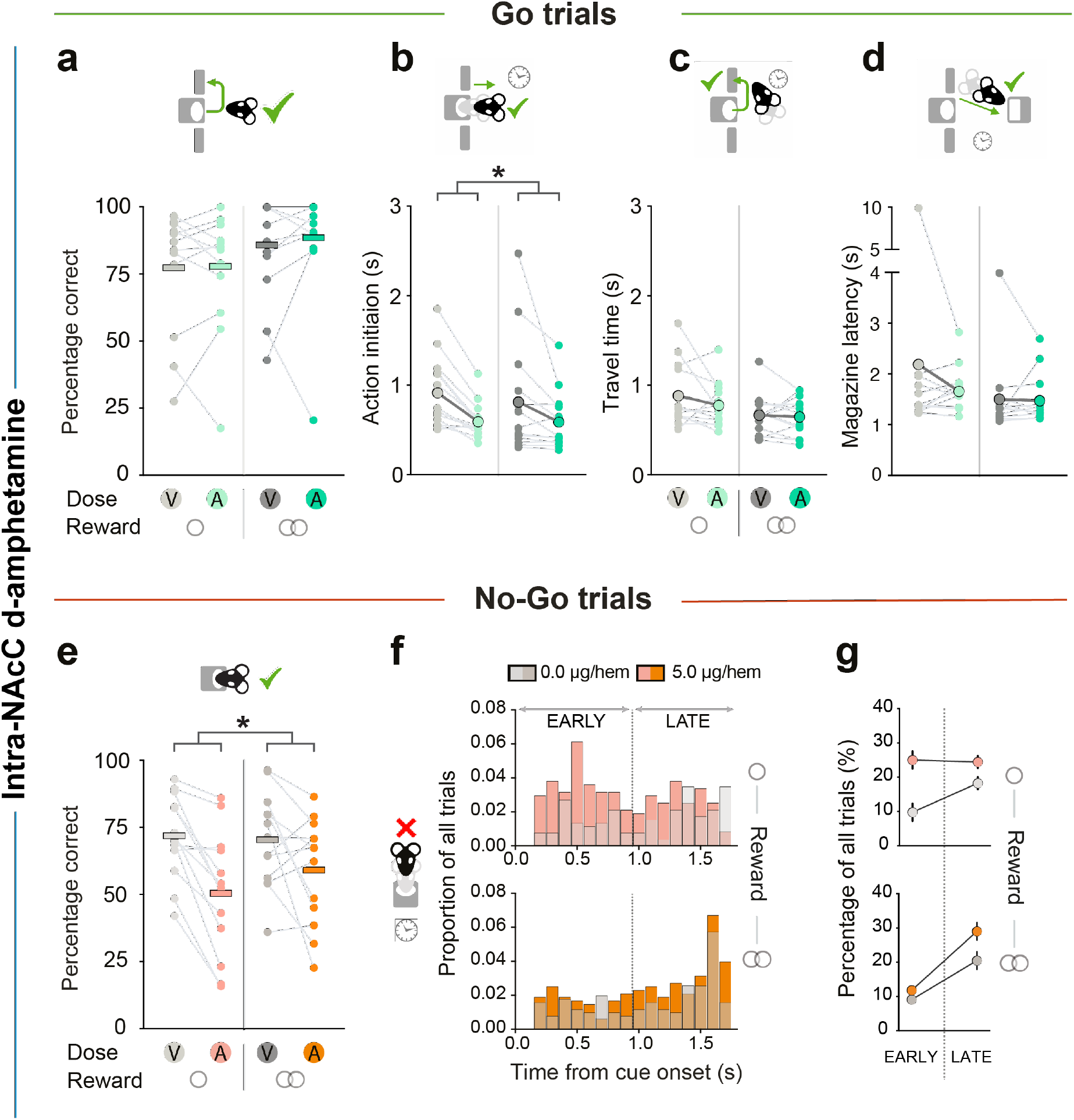
Effects of intra-NAcC d-amphetamine on Go/No-Go performance and latencies. **A.** Percentage of correct responses for small and large reward trials performed per drug session in Go trials. Data are shown as means (horizontal bars), superimposed by individual subject data (light grey lines and dots). Main effect of reward: F_1, 12_ = 5.37, p = 0.039. **B-D.** The effect of intra-NAcC d-amphetamine on action initiation, travel time and inter-press latency on Go trials. Data in B-D are depicted as means (large dots), superimposed by individual subject data (thin grey lines and small dots). Magazine latency: main effect of reward: F_1, 11_ = 8.51, p = 0.014. **E.** The effect of NAcC d-amphetamine on No-Go trial success rate. **F,G.** Timing of premature nose poke exits on unsuccessful small (upper panels) and large (lower panels) reward No-Go trials. Distributions are normalised to overall numbers of small or large reward No-Go trials. Data are shown in distributions of 100ms bins (F) or divided into ‘early’ and ‘late’ epochs of the No-Go hold requirement (G). Data are shown as mean ± SEM, normalized to the within-subject variance across the No-Go hold period of small (top panel) or large (bottom panel) reward trials. *p < .05.

On No-Go trials, intra-NAcC d-amphetamine markedly impaired rats’ ability to withhold action. It caused a substantial increase in No-Go errors, and these occurred *both* in the early and the late epochs of the No-Go holding interval (main effect of drug: F_1,12_=14.05 p=.003; all other F<3.66, p>.080) (fig.5e-g). Notably, these No-Go errors were often followed by lever pressing within the No-Go holding interval (F_1,12_=12.40, p=.004). This deficit in withholding actions also generalised to the pre-cue period, where intra-NAcC d-amphetamine substantially increased the numbers of aborted trials (main effect of drug: F_1,12_=5.76, p=.035; table S3).

Taken together, these findings demonstrate that intra-NAcC d-amphetamine significantly biased behavioural strategies towards action over inaction, speeding action initiation and impairing action restraint. In turn, this demonstrates that the null effects observed following intra-NAcC infusions of SB242084 cannot be simply attributed to tissue damage around the cannulae tips.

### Effect of SB242084 does not depend on training history or changes in task parameters

The systemic and local NAcC 5-HT_2C_ perturbations were carried out in separate cohorts, each having trained with slightly different No-Go hold intervals (1.7-1.9s and 1.5-1.7s for the systemic and the local 5-HT_2C_ manipulation, respectively). To ensure that the effects found were not attributable to any such differences in task parameters, or due to training experience, we analysed the effects of two replications of systemic administration of the low dose of SB242084 in the second cohort of animals, one administered before cannulae surgery and one performed after the infusion experiments had been completed.

Arguing against this possibility, systemic administration of SB242084 caused a very similar pattern of changes in task performance (see table 2). The ligand again improved Go trial success rate especially on small reward trials (drug X reward interaction: F_1,13_=13.56, p=.003, vehicle vs drug on small and large reward trials: p=.001 and p=.869, respectively). Likewise, the ligand again speeded up travel times and magazine latencies (main effect of drug: both F_1,13_>13.22, p<.004). Similarly, on No-Go trials, there was an interaction between the effects of the drug and reward, caused by a reduction in success rate in small but not large reward trials following SB242084 (drug X reward interaction: F_1,13_=7.14, p=.019). Importantly, there was no effect of whether the drug was given before or after the local infusion experiments in any of the above analyses (all F _1,13_<1.40, p>.258). Therefore, the lack of effects seen after the NAcC infusions of the 5-HT_2C_ ligand cannot simply be attributable to changes in specific task parameters or experience.

**Table 2.** Behavioural measures following two replications of administration of systemic SB242084 (Experiment 4). Data shown are the average across two experiments for each variable ± SEM (normalized to the within-subject variance across small and large reward trials). Asterisks and hats indicate statistical probability of a difference from the vehicle administration session(s) in *post hoc* tests.

|  | SMALL |  | LARGE |  |
| --- | --- | --- | --- | --- |
|  | vehicle | low dose | vehicle | low dose |
| Go success rate (%) | 82.28 ± 1.74 | 88.17 ± 1.35* | 93.94 ± 1.52 | 94.10 ± 1.46 |
| Response omission Go (%) | 13.93 ± 1.50 | 7.96 ± 1.64** | 4.46 ± 1.67 | 4.20 ± 1.41 |
| Wrong lever Go (%) | 3.78 ± 0.68 | 3.87 ± 0.53 | 1.53 ± 0.52 | 1.70 ± 0.61 |
| Go action initiation latency (s) | 0.75 ± 0.07 | 0.69 ± 0.06 | 0.65 ± 0.06 | 0.67 ± 0.06 |
| Go travel time (s) | 0.97 ± 0.03 | 0.83 ± 0.04** | 0.68 ± 0.04 | 0.57 ± 0.03** |
| Go food magazine latency (s) | 1.95 ± 0.12 | 1.92 ± 0.15* | 1.47 ± 0.13 | 1.39 ± 0.13* |
| No-Go success rate (%) | 73.89 ± 1.43 | 70.02 ± 1.62^ | 70.11 ± 1.43 | 71.40 ± 1.71 |

\* $p < .05$ , \*\* $p \leq .001$ , ^ $p=0.088$ , 2-tailed tests

## Discussion

Here we studied the role of 5-HT_2C_ receptors – an important modulator of instrumental vigour (Simpson et al. 2011; Bailey et al. 2016, 2018; Browne et al. 2017) – in controlling action and restraint. Subtle, yet consistent effects on both facets of behavioural control were detected. Systemic, but not intra-NAcC, administration of a low dose of a 5-HT_2C_ receptor antagonist SB242084, improved instrumental performance on Go trials. This was apparent even in the face of high baseline rates of accuracy, and was caused by a reduction in the rates of response omissions. Furthermore, although systemic 5-HT_2C_ antagonism had no effect on cued action initiation latencies, the drug dose-dependently speeded progress through the trial regardless of the reward size on offer. By contrast, 5-HT_2C_ blockade had a detrimental effect on goal-directed *restraint* of actions, but only on No-Go trials which promised small rewards. This was characterised by a potentiation of impulsive responses in the later part of the No-Go holding interval. This contrasted with the effects of intra-NAcC infusions of d-amphetamine which amplified *both* early and late premature action. Taken together, this suggests that 5-HT_2C_ receptors, outside the NAcC, play an important role in determining internally-driven response likelihood and instrumental vigour, shaped by the anticipated benefits of acting or restraint.

A number of previous studies, using a variety of tasks, have reported reduced operant responding latencies and increased motivation to work, particularly in high effort situations, following systemic administration of SB242084 (Simpson et al. 2011; Bailey et al. 2016, 2018; Browne et al. 2017). Here, we also observed increased success rate on Go trials and the dose-dependent invigoration of instrumental actions. Notably, however, this occurred in the context of a task with no equivalent effort requirements (note that while this task was arguably cognitively demanding, a recent study found no effect of 5-HT_2C_ receptor agents on cognitive effort allocation (Silveira et al. 2020). Therefore, while our results are generally consistent with the idea that perturbing transmission at 5-HT_2C_ receptors boosts goal-directed motivation and willingness to work (Simpson et al. 2011; Browne et al. 2017; Bailey et al. 2018), the current data refine these definitions and also suggest that there are potentially several distinct processes at play.

First, it was not the case that all response latencies were faster after systemic SB242084 as the average speed of cued action initiation on both Go and No-Go trials remained unchanged. This would suggest that the effect of the drug is not simply to boost motor output or attention, but is more specific in this task to the invigoration of ongoing reward-seeking movements. Second, while there was a monotonic effect with increasing drug dose on those latencies that were affected, the change in Go trial response omissions was limited to the low dose. This divergence between the effect of SB242084 on latencies and omissions is consistent with several previous reports (Winstanley et al. 2004b; Fletcher et al. 2007; Silveira et al. 2020). One possibility is that this reflects a shift in the balance between response speed and precision. While the low dose of the 5-HT_2C_ receptor antagonist enabled rats to respond faster *without* making them less likely to choose the correct lever or make more lever presses than required, the additional reduction in response times at the high dose might have started to make their responding more imprecise. Future studies, using a greater range of drug doses and video tracking of behaviour, will be needed to test this idea.

Third, there was some evidence that performance, though not response latencies, was most affected when the small reward was on offer. Although this might in part reflect a ceiling effect given high levels of baseline performance on large reward Go trials, note that a similar selective effect of the drug when the small reward was on offer was also observed for No-Go performance. Further, this is in line with previous work showing that the most prominent effects of SB242084 were apparent when animals were working for lower net value options (Bailey et al. 2016; Browne and Fletcher 2016). Therefore, the regulation of instrumental performance by 5-HT_2C_ receptors might depend on the opportunity cost – i.e., the magnitude of what would be lost by failing to perform appropriately. While the SB242084-driven increased engagement with low net value options may be advantageous in some circumstances, it can also impede flexible behaviour when the goal requires withholding a response.

In parallel to the improvements in Go trial performance, we also observed an increase in inappropriate, premature responses on No-Go trials following systemic SB242084, which again was specific to small reward trials. Moreover, there was evidence that the increase in No-Go failures was not reliably coupled to the increase in Go trial performance or responding vigour, indicating the changes are likely not attributable to the same psychological mechanism. There is much evidence from both manipulation (Harrison et al. 1997a, b) and physiological (Miyazaki et al. 2012; Fonseca et al. 2015) approaches that central serotonin is a key modulator of the ability to wait for reward, with 5-HT_2C_ receptors playing a central role in mediating this (Higgins et al. 2003, 2020; Winstanley et al. 2004b; Fletcher et al. 2007; Quarta et al. 2012; Silveira et al. 2020). However, the effect we observed here did not manifest as an overall increase in impulsivity nor a gross timing deficit, again suggesting that it is not simply a manifestation of a general increase in motor output. Instead, close inspection of the pattern of errors on these trials showed that the ligand specifically increased the proportion of errors that occurred in the *late* period of the No-Go holding interval on small reward trials, but had no influence on the rate of fast, cue-elicited No-Go errors. Moreover, when the animals were able correctly to withhold responding during the No-Go period, the 5-HT_2C_ receptor ligand shifted the pattern of subsequent action initiation latencies away from cue-elicited responses (i.e., ones clustered around cue offset at the end of the No-Go holding period or reward delivery 1s later).

Taken together with the absence of an observed effect on action initiation latencies, this points towards 5-HT_2C_ receptors having a more prominent influence over non-cue-driven processes, resulting in increased instrumental drive to act. Such an effect of SB242084 administration was also reported in a gambling task, where the ligand caused a selective amelioration of the influence of cues on risk-based decision-making (Adams et al. 2017). More generally, this perspective is compatible with recent evidence from optogenetic activation of dorsal raphe serotonin neurons that suggested serotonin may modulate the speed of evidence accumulation when deciding whether or not to switch away from a current behavioural policy, which in turn can also influence the vigour of the ongoing behaviour (Lottem et al. 2018). However, while the authors of this study suggested this was caused by serotonin modulating levels of uncertainty, it is unlikely that this factor is playing any major role here as the rats in the current study were highly trained and the cue-action-reward contingencies were deterministic and fixed. Instead, our data implicates 5-HT_2C_ receptors more closely with the regulation of instrumental drive based on the future benefits and opportunity costs of acting.

We had hypothesised that one potential locus for the effects of systemic administration of SB242084 on instrumental drive and restraint would be the NAcC. Not only is there a notable 5-HT projection to (Azmitia and Segal 1978; Vertes 1991) and level of expression of 5-HT_2C_ receptors in this region (Pazos et al. 1985; Eberle-Wang et al. 1997), but also a previous study, using the same doses of SB242084, reported an increase in impulsivity on the 5-choice serial reaction time task (5-CSRTT) specifically after infusions into NAcC and not into medial frontal regions (Robinson et al. 2008). Therefore, it was unexpected that intra-NAcC administration of SB242084 had no effect on any performance measure. This discrepancy was not caused by the cannulae targeting part of the NAcC that is not important for the task or from tissue damage caused by insertion of the injectors, as subsequent microinjections of *d*-amphetamine into the NAcC had a substantial influence on performance. Nor can the lack of effect result from subtle task or training differences in the different cohorts of animals that underwent the main systemic experiment and the NAcC cannulation respectively. This is because systemic administration of SB242084 in the cannulated cohort replicated the original patterns of results in the first cohort. Instead, what the findings presented here favour is a possible fractionation of 5-HT_2C_-modulated “waiting” impulsivity (Bari and Robbins 2013; Dalley and Robbins 2017): depending on whether the animals have to withhold a specific response until a cue is presented (as happens in the 5-CSRTT) or, as here, to withhold competing motor responses in the presence of a cue predicting future reward. These findings are also compatible with the possibility that systemic SB242084 effected the speeding of reward-guided responding through the dorsal striatum (Bailey et al. 2018) and inappropriate action through the medial prefrontal cortex (Miyazaki et al. 2020).

Previous studies have demonstrated a direct influence of systemic SB242084 on ventral tegmental area dopamine firing rates and dopamine levels in the NAcC (Matteo et al. 1999; De Deurwaerdere et al. 2004; Browne et al. 2017), and mesolimbic dopamine is known to shape goal-directed motivation and the balance of action initiation and restraint (Roitman et al. 2004; Wassum et al. 2012; Hamid et al. 2016; Syed et al. 2016; Dalley and Robbins 2017). Here, microinjections of *d*-amphetamine into NAcC, known to potentiate and prolong dopamine and, to a less extent, serotonin (Kankaanpää et al. 1998), also caused a marked increase in premature responses on No-Go trials and speeded action latencies on Go trials. However, the pattern of these changes was strikingly different to those observed after systemic administration of SB242084. Specifically, d-amphetamine exclusively speeded action initiation latencies on Go trials, which had been unaffected by administration of SB242084, but had no effect on the speed of other responses in a trial or overall success rates, all of which had been altered by systemic 5-HT_2C_ receptor manipulations. Similarly, premature responses were elevated both in the pre-cue period and throughout the No-Go holding interval after NAcC d-amphetamine, as compared to a selective elevation in the later No-Go holding period after systemic SB242082. This raises the interesting possibility that mesolimbic dopamine and serotonin, acting through 5-HT_2C_Rs, provide complementary but distinct modulation of instrumental drive and response restraint, with the former regulating the rapid initiation of reward-seeking actions and the latter affecting instrumental drive for reward. Future studies where the two systems are directly manipulated in the context of the same behavioural task may help to further substantiate the existence of these complementary roles. Note though that this does not rule out additional, more direct interactions between 5-HT_2C_ transmission and dopamine elsewhere in the basal ganglia implicated in regulating action restraint and response vigour. For example, Bailey and colleagues demonstrated dorsal striatal dopamine levels to be a crucial modulator of instrumental performance after administration of SB242084 (Eberle-Wang et al. 1997; Agnoli and Carli 2012; Bailey et al. 2018). Furthermore, SB242084 could be acting in a dopamine-independent manner in the dorsal striatum or elsewhere in the basal ganglia such as the subthalamic nucleus or NAc shell (Eberle-Wang et al. 1997; Filip and Cunningham 2002; Agnoli and Carli 2012; Bailey et al. 2018).

The effect of SB242084 on goal-directed instrumental vigour has led to the possibility that ligands targeting the 5-HT_2C_R – and in particular SB242084 given its functional selectivity over signalling pathways coupled to 5-HT_2C_ receptors – could potentially be used to treat patients with motivational deficits such as apathy (Matteo et al. 1999; Simpson et al. 2011; Bailey et al. 2018). Our and others’ data add a note of caution by showing that the improvement in instrumental drive for reward may, in certain contexts, also have detrimental effects on response restraint. Similarly, 5-HT_2C_R blockade has been observed both to reduce and to amplify particular behaviours associated with obsessive-compulsive disorders (Martin et al. 2002; Albelda and Joel 2012). Therefore, further research will be required to understand the neural mechanisms underlying the complex changes in response vigour and in response restraint reported here, to allow for more precise targeting of 5-HT_2C_R and its associated signalling pathways in the future.

## Acknowledgements

We would like to thank Sanjay Manohar and Sean James Fallon for analysis advice, and Greg Daubney for help with histology.

## References

Adams WK, Barkus C, Ferland JMN, et al (2017) Pharmacological evidence that 5-HT2C receptor blockade selectively improves decision making when rewards are paired with audiovisual cues in a rat gambling task. Psychopharmacology (Berl) 234:3091–3104. https://doi.org/10.1007/s00213-017-4696-4

Agnoli L, Carli M (2012) Dorsal-striatal 5-HT 2A and 5-HT 2C receptors control impulsivity and perseverative responding in the 5-choice serial reaction Time Task. Psychopharmacology (Berl) 219:633–645. https://doi.org/10.1007/s00213-011-2581-0

Albelda N, Joel D (2012) Current animal models of obsessive compulsive disorder: An update. Neuroscience 211:83–106. https://doi.org/10.1016/j.neuroscience.2011.08.070

Azmitia EC, Segal M (1978) An autoradiographic analysis of the differential ascending projections of the dorsal and median raphe nuclei in the rat. J Comp Neurol 179:641–667. https://doi.org/10.1002/cne.901790311

Bailey MR, Goldman O, Bello EP, et al (2018) An Interaction between Serotonin Receptor Signaling and Dopamine Enhances Goal-Directed Vigor and Persistence in Mice. J Neurosci 38:2088–17. https://doi.org/10.1523/JNEUROSCI.2088-17.2018

Bailey MR, Williamson C, Mezias C, et al (2016) The effects of pharmacological modulation of the serotonin 2C receptor on goal-directed behavior in mice. Psychopharmacology (Berl) 233:615–624. https://doi.org/10.1007/s00213-015-4135-3

Bari A, Robbins TW (2013) Inhibition and impulsivity: Behavioral and neural basis of response control. Prog Neurobiol 108:44–79. https://doi.org/10.1016/j.pneurobio.2013.06.005

Browne CJ, Fletcher PJ (2016) Decreased Incentive Motivation Following Knockout or Acute Blockade of the Serotonin Transporter: Role of the 5-HT2C Receptor. Neuropsychopharmacology 41:2566–2576. https://doi.org/10.1038/npp.2016.63

Browne CJ, Ji X, Higgins GA, et al (2017) Pharmacological Modulation of 5-HT 2C Receptor Activity Produces Bidirectional Changes in Locomotor Activity, Responding for a Conditioned Reinforcer, and Mesolimbic DA Release in C57BL /6 Mice. Neuropsychopharmacology 42:2178–2187. https://doi.org/10.1038/npp.2017.124

Cohen JY, Amoroso MW, Uchida N (2015) Serotonergic neurons signal reward and punishment on multiple timescales. Elife 4:1–25. https://doi.org/10.7554/eLife.06346

Cole B, Robbins T (1987) Amphetamine impairs the discriminative performance of rats with dorsal noradrenergic bundle lesions on a 5-choice serial reaction time task: new evidence for central dopaminergic-noradrenergic interactions. Psychopharmacology (Berl) 91:458–66. https://doi.org/10.1007/BF00216011

Cools R, Nakamura K, Daw ND (2011) Serotonin and dopamine: unifying affective, activational, and decision functions. Neuropsychopharmacology 36:98–113. https://doi.org/10.1038/npp.2010.121

Dalley JW, Robbins TW (2017) Fractionating impulsivity: Neuropsychiatric implications. Nat Rev Neurosci 18:158–171. https://doi.org/10.1038/nrn.2017.8

De Deurwaerdère P, Lagière M, Bosc M, Navailles S (2013) Multiple controls exerted by 5-HT2C receptors upon basal ganglia function: From physiology to pathophysiology. Exp Brain Res 230:477–511. https://doi.org/10.1007/s00221-013-3508-2

De Deurwaerdere P, Navailles S, Berg KA, et al (2004) Constitutive Activity of the Serotonin2C Receptor Inhibits In Vivo Dopamine Release in the Rat Striatum and Nucleus Accumbens. J Neurosci 24:3235–3241. https://doi.org/10.1523/JNEUROSCI.0112-04.2004

Di Matteo V, Di Giovanni G, Esposito E (2000) SB 242084: A selective 5-HT(2C) receptor antagonist. CNS Drug Rev 6:195–205. https://doi.org/10.1111/j.1527-3458.2000.tb00147.x

du Hoffmann J, Nicola SM (2014) Dopamine invigorates reward seeking by promoting cue-evoked excitation in the nucleus accumbens. J Neurosci 34:14349–14364. https://doi.org/10.1523/JNEUROSCI.3492-14.2014

Dudman JT, Krakauer JW (2016) The basal ganglia: From motor commands to the control of vigor. Curr Opin Neurobiol 37:158–166. https://doi.org/10.1016/j.conb.2016.02.005

Eberle-Wang K, Mikeladze Z, Uryu K, Chesselet MF (1997) Pattern of expression of the serotonin 2C receptor messenger RNA in the basal ganglia of adult rats. J Comp Neurol 384:233–247

Filip M, Cunningham KA (2002) Serotonin 5-HT2C receptors in nucleus accumbens regulate expression of the hyperlocomotive and discriminative stimulus effects of cocaine. Pharmacol Biochem Behav 71:745–756. https://doi.org/10.1016/S0091-3057(01)00741-9

Fletcher PJ, Tampakeras M, Sinyard J, Higgins GA (2007) Opposing effects of 5-HT2A and 5-HT2C receptor antagonists in the rat and mouse on premature responding in the five-choice serial reaction time test. Psychopharmacology (Berl) 195:223–234. https://doi.org/10.1007/s00213-007-0891-z

Floresco SB (2015) The nucleus accumbens: An interface between cognition, emotion, and action. Annu Rev Psychol 66:25–32. https://doi.org/10.1146/annurev-psych-010213-115159

Fonseca MS, Murakami M, Mainen ZF (2015) Activation of dorsal raphe serotonergic neurons promotes waiting but is not reinforcing. Curr Biol 25:306–315. https://doi.org/10.1016/j.cub.2014.12.002

Grima L, Panayi M, Haermson O, et al (2021) Nucleus accumbens D1-receptors regulate and focus transitions to reward-seeking action 2. bioRxiv 2021.06.15.448563

Hamid AA, Pettibone JR, Mabrouk OS, et al (2016) Mesolimbic dopamine signals the value of work. Nat Neurosci 19:117–126. https://doi.org/10.1038/nn.4173

Harrison AA, Everitt BJ, Robbins TW (1997a) Central 5-HT depletion enhances impulsive responding without affecting the accuracy of attentional performance: Interactions with dopaminergic mechanisms. Psychopharmacology (Berl) 133:329–342. https://doi.org/10.1007/s002130050410

Harrison AA, Everitt BJ, Robbins TW (1997b) Doubly dissociable effects of median- and dorsal-raphe lesions on the performance of the five-choice serial reaction time test of attention in rats. Behav Brain Res 89:135–149. https://doi.org/10.1016/S0166-4328(97)00053-3

Higgins GA, Brown M, St John J, et al (2020) Effects of 5-HT2C receptor modulation and the NA reuptake inhibitor atomoxetine in tests of compulsive and impulsive behaviour. Neuropharmacology 170:108064. https://doi.org/10.1016/j.neuropharm.2020.108064

Higgins GA, Enderlin M, Haman M, Fletcher PJ (2003) The 5-HT2A receptor antagonist M100,907 attenuates motor and “impulsive-type” behaviours produced by NMDA receptor antagonism. Psychopharmacology (Berl) 170:309–319. https://doi.org/10.1007/s00213-003-1549-0

Higgins GA, Fletcher PJ (2003) Serotonin and drug reward: Focus on 5-HT2C receptors. Eur J Pharmacol 480:151–162. https://doi.org/10.1016/j.ejphar.2003.08.102

Kankaanpää A, Meririnne E, Lillsunde P, Seppälä T (1998) The acute effects of amphetamine derivatives on extracellular serotonin and dopamine levels in rat nucleus accumbens. Pharmacol Biochem Behav 59:1003–1009. https://doi.org/10.1016/S0091-3057(97)00527-3

Kennett GA, Wood MD, Bright F, et al (1997) SB 242084, a selective and brain penetrant 5-HT(2C) receptor antagonist. Neuropharmacology 36:609–620. https://doi.org/10.1016/S0028-3908(97)00038-5

Ko D, Wanat MJ (2016) Phasic dopamine transmission reflects initiation vigor and exerted effort in an action- and region-specific manner. J Neurosci 36:2202–2211. https://doi.org/10.1523/JNEUROSCI.1279-15.2016

Kulichenko AM, Pavlenko VB (2004) Self-Initiated Behavioral Act-Related Neuronal Activity in the Region of the Raphe Nuclei of the Cat. Neurophysiology 36:50–57. https://doi.org/10.1023/B:NEPH.0000035969.92198.9f

Li Y, Zhong W, Wang D, et al (2016) Serotonin neurons in the dorsal raphe nucleus encode reward signals. Nat Commun 7:10503. https://doi.org/10.1038/ncomms10503

Lottem E, Banerjee D, Vertechi P, et al (2018) Activation of serotonin neurons promotes active persistence in a probabilistic foraging task. Nat Commun 9:1000. https://doi.org/10.1038/s41467-018-03438-y

Martin JR, Ballard TM, Higgins GA (2002) Influence of the 5-HT2C receptor antagonist, SB-242084, in tests of anxiety. Pharmacol Biochem Behav 71:615–625. https://doi.org/10.1016/S0091-3057(01)00713-4

Matias S, Lottem E, Dugue GP, Mainen ZF (2017) Activity patterns of serotonin neurons underlying cognitive flexibility. Elife 6:1–24. https://doi.org/10.7554/eLife.20552

Matteo V Di, Giovanni G Di, Mascio M Di, Esposito E (1999) SB 242 084, a selective serotonin 2C receptor antagonist, increases dopaminergic transmission in the mesolimbic system. Neuropharmacology 38:1195–1205. https://doi.org/10.1016/s0028-3908(99)00047-7

Mcelroy JF, Dupont AF, Feldman RS (1982) The Effects of Fenfluramine and Fluoxetine on the Acquisition of a Conditioned Avoidance Response in Rats. Psychopharmacology (Berl) 77:356–359. https://doi.org/10.1007/BF00432770

Miyazaki K, Miyazaki KW, Doya K (2011) Activation of dorsal raphe serotonin neurons underlies waiting for delayed rewards. J Neurosci 31:469–479. https://doi.org/10.1523/JNEUROSCI.3714-10.2011

Miyazaki K, Miyazaki KW, Doya K (2012) The role of serotonin in the regulation of patience and impulsivity. Mol Neurobiol 45:213–224. https://doi.org/10.1007/s12035-012-8232-6

Miyazaki K, Miyazaki KW, Sivori G, et al (2020) Serotonergic projections to the orbitofrontal and medial prefrontal cortices differentially modulate waiting for future rewards. Sci Adv 6:1–15. https://doi.org/10.1126/sciadv.abc7246

Pattij T, Janssen MCW, Vanderschuren LJMJ, et al (2007) Involvement of dopamine D1 and D2 receptors in the nucleus accumbens core and shell in inhibitory response control. Psychopharmacology (Berl) 191:587–598. https://doi.org/10.1007/s00213-006-0533-x

Pazos A, Cortés R, Palacios JM (1985) Quantitative autoradiographic mapping of serotonin receptors in the rat brain. II. Serotonin-2 receptors. Brain Res 346:231–249. https://doi.org/10.1016/0006-8993(85)90856-X

Petzold GC, Hagiwara A, Murthy VN (2009) Serotonergic modulation of odor input to the mammalian olfactory bulb. Nat Neurosci 12:784–791. https://doi.org/10.1038/nn.2335

Pothuizen HHJ, Jongen-Rêlo AL, Feldon J, Yee BK (2005) Double dissociation of the effects of selective nucleus accumbens core and shell lesions on impulsive-choice behaviour and salience learning in rats. Eur J Neurosci 22:2605–2616. https://doi.org/10.1111/j.1460-9568.2005.04388.x

Quarta D, Naylor CG, Glennon JC, Stolerman IP (2012) Serotonin antagonists in the five-choice serial reaction time task and their interactions with nicotine. Behav Pharmacol 23:143–152. https://doi.org/10.1097/FBP.0b013e32834f9fb4

Robbins TW, Everitt BJ (1992) Functions of dopamine in the dorsal and ventral striatum. Semin Neurosci 4:119–127. https://doi.org/10.1016/1044-5765(92)90010-Y

Robinson ESJ, Dalley JW, Theobald DEH, et al (2008) Opposing Roles for 5-HT 2A and 5-HT 2C Receptors in the Nucleus Accumbens on Inhibitory Response Control in the 5-Choice Serial Reaction Time Task. Neuropsychopharmacology 33:2398–2406. https://doi.org/10.1038/sj.npp.1301636

Roitman MF, Stuber GD, Phillips PEM, et al (2004) Dopamine Operates as a Subsecond Modulator of Food Seeking. J Neurosci 24:1265–1271. https://doi.org/10.1523/JNEUROSCI.3823-03.2004

Salamone JD, Correa M, Farrar A, Mingote SM (2007) Effort-related functions of nucleus accumbens dopamine and associated forebrain circuits. Psychopharmacology (Berl) 191:461–482. https://doi.org/10.1007/s00213-006-0668-9

Silveira MM, Wittekindt SN, Mortazavi L, et al (2020) Investigating serotonergic contributions to cognitive effort allocation, attention, and impulsive action in female rats. J Psychopharmacol 34:452–466. https://doi.org/10.1177/0269881119896043

Simpson EH, Kellendonk C, Ward RD, et al (2011) Pharmacologic rescue of motivational deficit in an animal model of the negative symptoms of schizophrenia. Biol Psychiatry 69:928–935. https://doi.org/10.1016/j.biopsych.2011.01.012

Soubrié P (1986) Reconciling the role of central serotonin neurons in human and animal behavior. Behav Brain Sci 9:319. https://doi.org/10.1017/S0140525X00022871

Syed ECJ, Grima LL, Magill PJ, et al (2016) Action initiation shapes mesolimbic dopamine encoding of future rewards. Nat Neurosci 19:1–6. https://doi.org/10.1038/nn.4187

Vertes RP (1991) A PHA-L analysis of ascending projections of the dorsal raphe nucleus in the rat. J Comp Neurol 313:643–668. https://doi.org/10.1002/cne.903130409

Wassum KM, Ostlund SB, Maidment NT (2012) Phasic mesolimbic dopamine signaling precedes and predicts performance of a self-initiated action sequence task. Biol Psychiatry 71:846–854. https://doi.org/10.1016/j.biopsych.2011.12.019

West CHK, Boss-Williams KA, Weiss JM (1998) Motor activation by amphetamine infusion into nucleus accumbens core and shell subregions of rats differentially sensitive to dopaminergic drugs. Behav Brain Res 98:155–165. https://doi.org/10.1016/S0166-4328(98)00064-3

Winstanley CA, Dalley JW, Theobald DEH, Robbins TW (2004a) Fractionating impulsivity: contrasting effects of central 5-HT depletion on different measures of impulsive behavior. Neuropsychopharmacology 29:1331–1343. https://doi.org/10.1038/sj.npp.1300434

Winstanley CA, Theobald DEH, Dalley JW, et al (2004b) 5-HT2A and 5-HT2C receptor antagonists have opposing effects on a measure of impulsivity: Interactions with global 5-HT depletion. Psychopharmacology (Berl) 176:376–385. https://doi.org/10.1007/s00213-004-1884-9

