## Supplemental Material for "5-HT_2C_ receptor perturbation has bidirectional influence over instrumental vigour and restraint"

**Supplementary table 1.** Additional behavioural measures following systemic SB242084 (Experiment 1). Data shown are the mean for each variable (± SEM, normalized to the within-subject variance across small and large reward trials). Asterisks indicate significant difference from the vehicle administration session(s) in *post hoc* tests.

|  | SMALL | | | LARGE | | |
| --- | --- | --- | --- | --- | --- | --- |
|  | vehicle | low dose | high dose | vehicle | low dose | high dose |
| Go wrong lever choice (%) | 3.10 ± 0.84 | 2.82 ± 1.07 | 4.89 ± 1.32 | 1.71 ± 0.83 | 1.40 ± 0.50 | 2.43 ± 1.21 |
| Aborted trials per trial | 0.29 ± 0.04 | 0.24 ± 0.02 | 0.21 ± 0.04 | 0.33 ± 0.05 | 0.32 ± 0.03 | 0.32 ± 0.04 |
| Go Lever presses | 2.85 ± 0.14 | 2.96 ± 0.16 | 2.89 ± 0.13 | 2.88 ± 0.19 | 3.13 ± 0.16 | 2.91 ± 0.11 |
| Go Inter-press latency (s) | 0.72 ± 0.04 | 0.78 ± 0.10 | 0.64 ± 0.05* | 0.57 ± 0.06 | 0.49 ± 0.06 | 0.46 ± 0.06* |
| No-Go success rate (%) | 83.99 ± 2.56 | 73.11 ± 2.16* | 76.68 ± 2.50* | 86.00 ± 1.79 | 84.44 ± 2.14 | 85.60 ± 2.31 |
| No-Go magazine latency (s) | 2.18 ± 0.1 | 2.04 ± 0.08* | 1.93 ± 0.07* | 1.71 ± 0.11 | 1.75 ± 0.07* | 1.69 ± 0.08* |
| Re-engagement latency (s) | 4.23 ± 0.40 | 3.60 ± 0.27 | 3.72 ± 0.19 | 3.97 ± 0.28 | 3.28 ± 0.22 | 3.47 ± 0.20 |

*p < .05, **p < .001

**Supplementary table 2.** Additional behavioural measures following intra-NAcC SB242084 (Experiment 2). Data shown are the mean for each variable (± SEM, normalized to the within-subject variance across small and large reward trials). Asterisks indicate significant difference from the vehicle administration session(s) in *post hoc* tests.

|  | SMALL | | | LARGE | | |
| --- | --- | --- | --- | --- | --- | --- |
|  | vehicle | low dose | high dose | vehicle | low dose | high dose |
| Go wrong lever choice (%) | 5.40 ± 0.67 | 5.83 ± 1.72 | 4.38 ± 1.38 | 5.56 ± 0.80 | 4.50 ± 0.56 | 5.47 ± 2.01 |
| Aborted trials per trial | 0.39 ± 0.05 | 0.32 ± 0.04 | 0.36 ± 0.08 | 0.52 ± 0.04 | 0.47 ± 0.08 | 0.58 ± 0.06 |
| Go Lever presses | 3.03 ± 0.10 | 3.14 ± 0.14 | 3.25 ± 0.14 | 3.11 ± 0.16 | 2.81 ± 0.18 | 2.80 ± 0.11 |
| Go Inter-press latency (s) | 0.64 ± 0.10 | 0.68 ± 0.70 | 0.66 ± 0.08 | 0.62 ± 0.09 | 0.66 ± 0.10 | 0.77 ± 0.07 |
| No-Go magazine latency (s) | 2.59 ± 0.05 | 2.48 ± 0.06 | 2.65 ± 0.06 | 2.43 ± 0.04 | 2.36 ± 0.04 | 2.56 ± 0.06 |
| Re-engagement latency (s) | 3.52 ± 0.26 | 3.32 ± 0.23 | 3.20 ± 0.14 | 3.83 ± 0.18 | 3.32 ± 0.22 | 3.35 ± 0.19 |

**Supplementary table 3.** Additional behavioural measures following intra-NAcC d-amphetamine (Experiment 3). Data shown are the mean for each variable (± SEM, normalized to the within-subject variance across small and large reward trials). Asterisks indicate significant difference from the vehicle administration session(s) in *post hoc* tests.

|  | SMALL | | LARGE | |
| --- | --- | --- | --- | --- |
|  | vehicle | d-amphetamine | vehicle | d-amphetamine |
| Go wrong lever choice (%) | 7.48 ± 1.51 | 8.74 ± 1.39 | 3.33 ± 1.33 | 3.68 ± 0.62 |
| Aborted trials per trial | 0.36± 0.05 | 0.73 ± 0.09* | 0.42 ± 0.08 | 0.69 ± 0.05* |
| Go Lever presses | 2.84 ± 0.12 | 2.84 ± 0.18 | 2.76 ± 0.18 | 2.77 ± 0.16 |
| Go Inter-press latency (s) | 0.66 ± 0.05 | 0.64 ± 0.06 | 0.59 ± 0.08 | 0.62 ± 0.05 |
| No-Go magazine latency (s) | 2.60 ± 0.08 | 2.31 ± 0.06 | 2.39 ± 0.04 | 2.26 ± 0.06 |
| Re-engagement latency (s) | 3.60 ± 0.20 | 3.60 ± 0.21 | 3.45 ± 0.21 | 3.19 ± 0.18 |

*p < .05, **p < .001
